# Graph–based integration of histone modifications profiles: haematopoietic cell differentiation as a case study

**DOI:** 10.1101/2020.10.22.350611

**Authors:** Federica Baccini, Monica Bianchini, Filippo Geraci

## Abstract

In this paper, we show that quantifying histone modifications by counting the number of high– resolution peaks in each gene allows to build profiles of these epigenetic marks, associating them to a phenotype. The significance of this approach is verified by applying graph–cut techniques for assessing the differentiation between myeloid and lymphoid cells in haematopoiesis, i.e. the process through which all the different types of blood cells originate starting from a unique cell type. The experiments are conducted on a population of samples from 24 cell types involved in haematopoiesis. Six profiles are constructed for each cell type, based on a different histone modification signal. Following the experimentally verified idea that the peak number distribution per gene behaves similarly to gene expression, the profile computation employs standard differential analysis tools to find genes whose epigenetic modifications are related to a given phenotype. Next, six similarity networks of cell types are constructed, based on each histone modification, and then combined into a unique one through similarity network fusion. Finally, the similarity networks are transformed into dissimilarity graphs, to which two different cuts are applied and compared to evaluate the classic differentiation between myeloid and lymphoid cells. The results show that all histone modifications contribute almost equally to the myeloid/lymphoid differentiation, and this is also confirmed by the analysis of the fused network. However, they also suggest that histone modifications may not be the only mechanism for regulating the differentiation of hematopoietic cells.

## 1 Introduction

A histone modification is a covalent post–translational modification to histone proteins, which includes methylation, phosphorylation, acetylation, ubiquitylation, and sumoylation. Histone modifications are epigenetic marks known to modify the structure of chromatin, with the effect of regulating gene expression [1]. In particular, histone acetylation [5] is a dynamic process regulated by two family of enzymes, namely histone acetyltransferases (HATs) and deacetylases (HDACs). HATs use coenzyme A (CoA) as cofactor to catalyze the transfer of an acetyl group to the amine group of a lysine side chain. This reaction produces a neutralization of the positively charged lysine, leading to a weakening of the interaction between histones and DNA (which is negatively charged). HDAC has an opposite effect with respect to HAT, in that it reverses lysine acetylation, thus restoring its positive charge. Therefore, it has a stabilyzing action, which is consistent with the fact that it is mainly involved in transcriptional repression. Methylation, instead, is a type of modification occuring mainly on the side chains of lysines and arginines. It consists in the addition of one, two or three methyl groups to the side chain of the interested aminoacid (mono–, di–, or tri–methylation). The families of enzimes that are responsible for this modification are methylases and methyltransferases. The peculiarity of methylation is that it does not alter the histone charge.

It is known that different types of histone modifications can have either an inhibitory or promoting effect on gene transcription [22]. For instance, H3K27ac, and the mono and tri-–methylation of lysine 4 (H3K4me1 and H3K4me3) are often associated with promoters and enhancers, while tri-–methylation of lysine 27 (H3K27me3) and 36 (H3K36me3) are associated with transcriptional repression [8]. The role of H3K9me3 has yet to be understood. The variation among the potential interaction of marks and effectors suggests that there is an overall competition for modifications to achieve the proper chromatin state [6]. Therefore, it is relevant to study relations and interactions between different modifications in order to discover new regulatory patterns.

In this work, histone modifications are analysed in order to verify a hypothesis on haematopoiesis. Haematopoiesis is the differentiation process that begins from haematopoietic stem cells (HSCs) and leads to all the blood cell types through a series of differentiation steps [7]. It is a process that begins well before birth and continues throughout an individual’s life. In the embryo, the *primitive haematopoiesis* process produces only red blood cells, capable of supplying oxygen to developing organs. At this stage, indeed, the amniotic sac, which feeds the embryo until the placenta is fully developed, controls haematopoiesis. As the embryo continues to grow, the haematopoiesis process moves to the liver, spleen and bone marrow and begins to produce other types of blood cells. In infants and children, haematopoiesis of red blood cells and platelets occurs in the bone marrow, but it can also continue in the spleen and liver. In adults, haematopoiesis of red blood cells and platelets occurs mainly in the bone marrow. The lymphatic system, especially the spleen, lymph nodes and thymus, produce a type of white blood cell called *lymphocyte*, while *monocytes* are released mainly by the bone marrow. The body continually produces new red and white blood cells to replace old ones. About 1% of the body’s blood cells need to be replaced every day, though the haematopoiesis rate depends on the needs of the body. White blood cells have a shorter average–life, from a few hours to a few days, while red blood cells can last up to about 120 days.

In the classic model of haematopoiesis, the different blood cell lines originate from a process that follows a hierarchical pattern, in which the differentiation potential gradually decreases (see Figure 1). The starting point is represented by multipotent cells, from which oligopotent cells originate, each able to differentiate only in a few cell lines. Finally, unipotent cells are obtained, which can originate only one type of cell. Until recently, a simple model based on a series of binary branches — where the first two branches account for the *myeloid* and the *lymphoid* lineages — was broadly agreed, but with the advent of the single cell sequencing this model has shown its limits, raising the need of more articulated models [4]. In particular, in [14], the CD34+ multipotent haematopoietic cells and their progressive differentiation scheme have been studied in the different phases of the development of the human individual, from the fetus to adulthood. Such a differentiation scheme primarily gives rise to myeloid progenitors, which give rise to a significant part of the blood cells, including erythrocytes, commonly called red blood cells, or megakaryocytes, responsible for the platelet production. Indeed, if the liver of the fetus — where haematopoiesis takes place for a few weeks during gestation — contains a large number of distinct oligopotent progenitors, including myeloid cells, erythrocytes and megakaryocytes, in the bone marrow of the adult individual, the number of oligopotent progenitor cells is very limited. In summary, this means that, with adulthood, there is a transition to a two– level hierarchy, not foreseen in the current dominant model of haematopoiesis, developed since the 1960s, with different types of blood cells forming quickly from stem cells and not at the end of the entire hierarchical differentiation process. This discovery allowed to better understand a wide range of hematological diseases, from anemia, i.e. a pathological reduction of hemoglobin in the blood, to leukemia, which causes the uncontrolled proliferation of blood cells.

**Figure 1:**
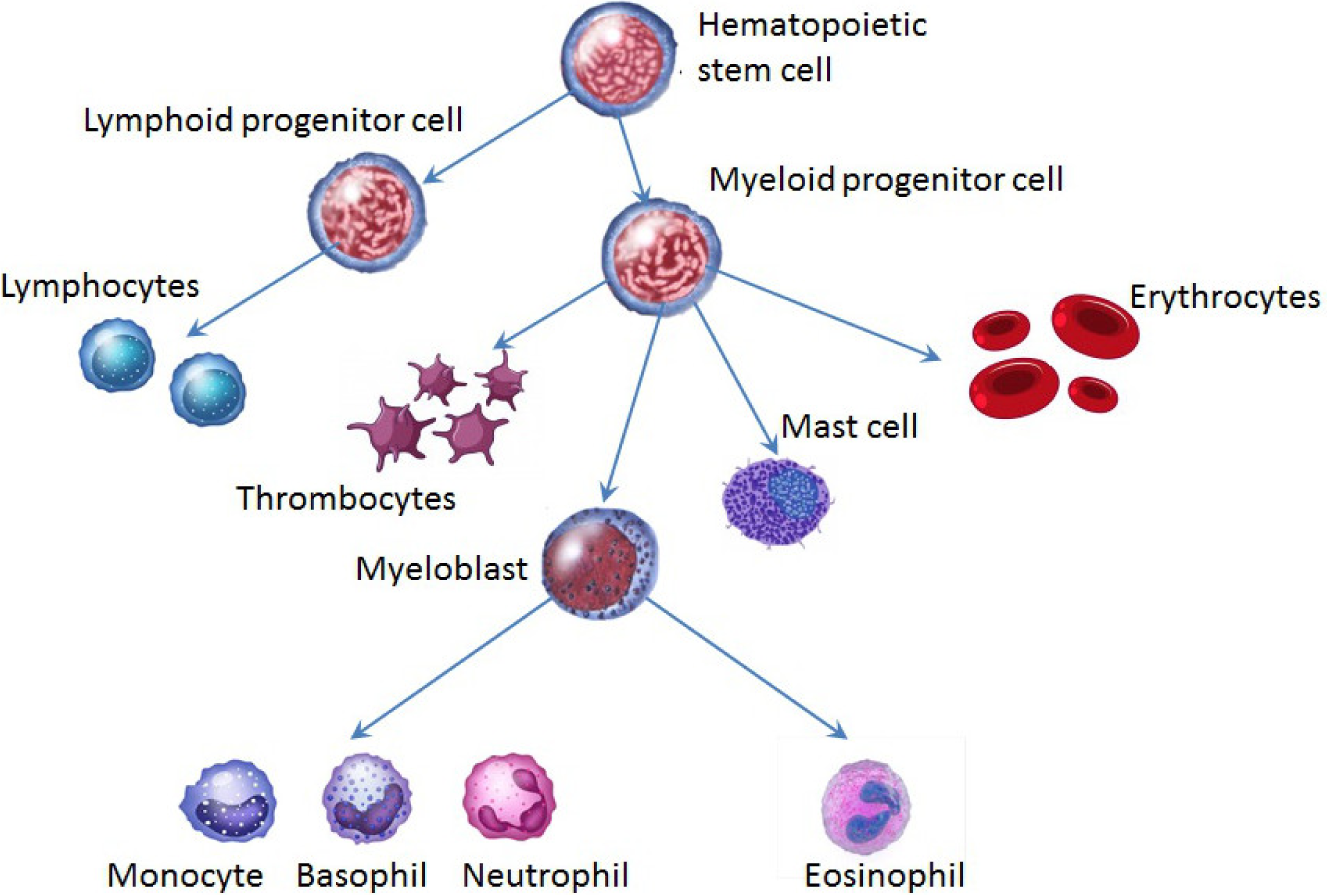
A simplified classical model of the haematopoietic tree.

In this paper, we introduce a quantitative method for verifying the plausibility of the differentiation between myeloid and lymphoid lineages, which characterizes the first hierarchical subdivision in the haematopoietic tree. First of all, networks of cells characterized by similar histone modification profiles are constructed. Then, a Similarity Network Fusion (SNF) algorithm [21] was applied to integrate them into a unique similarity network, through a cross diffusion process. The fused similarity network can be viewed as a way for synthesizing significant features coming from different data types or, in other words, as a viable solution for collecting information on how multiple variables determine similarities among entities constituting the network. Lastly, the goodness of the myeloid/lymphoid hypothesis has been evaluated by applying a graph– cut approach to networks of dissimilarity among cell types (represented by pairwise dissimilarity matrices). According to our model, the weight of the edge between two cell types is proportional to their dissimilarity. Consequently, if a given hypothesis is supported by data, partitioning the network on this basis will cause the removal of high weight edges. In fact, in order to evaluate the plausibility of a binary classification, each node of the dissimilarity network is labeled according to the class it belongs to. Then, the problem of verifying how good the distinction between the two classes is reduces to quantifying the cost of bi–partitioning the network into two sets, corresponding to the classes, and comparing such cost with that of the best cut. The experimental results agree with the classical myeloid/lymphoid hypothesis, although they suggest that this model cannot be the only mechanism for haematopoietic blood differentiation. Finally, and most interestingly, the approach proposed in this paper, tested on blood cell differentiation, can be applied to different problems in which the correctness of a particular hypothesis, supported by data, must be evaluated.

## 2 Method

Histone modification profiles from the IHEC database come as continuous signals, one for each of the 6 histone modifications marks identified by the Roadmap Epigenome Mapping Centers^1^, where each nucleotide is associated a quantification of the corresponding modifier. The profiles have been obtained by means of the ChIP–seq technology, which combines chromatin immunoprecipitation (ChIP) with DNA sequencing. The ChIP–seq protocol [15] exploits the possibility of establishing a covalent crosslink between histone proteins and the underlying DNA, in order to identify genomic regions where a histone modification is present. After crosslinking, indeed, the resulting protein– DNA complexes can be captured using histone–specific antibodies. Subsequently, the link is broken and the DNA is purified, enriched, and sequenced. This procedure causes the amount of DNA of a specific location to be proportional to the amount of a histone modification. As a result, quantification can be done by counting the number of reads piling up on each genomic location.

According to our hypothesis, DNA accessibility, and, in turn, gene expression ability, may depend not only on the presence/absence of acetylated/methylated histone sites, but also on the concentration of these modifications at certain locations, (i.e. on the formation of peaks). Indeed, the formation of such peaks may trigger noticeable modifications of the chromatin structure.

### 2.1 Peak calling

Let {*X* = *x*_1_, …, *x*_*n*_} be the histone modification profile of the sample cell *X*, where the value *x*_*i*_ corresponds to the number of supporting reads covering the genomic position *i*. Intuitively, a peak is a local maximum in *X*, i.e. a point whose value is reasonably higher than those in its surrounding. Instead of just considering the single maximum point, we can extend the definition of peak by including also the two monotone curves leading to the maximum. Despite simple in principle, the two free parameters (peak height and surrounding size) involved in the definition make finding peaks complicated, since different algorithms for specific needs may be required. Setting large surrounding areas leads to large peaks which are suitable for the identification of genomic sites involved in a histone modification (this is, for example, the case of Sole–Search [3] — the peak detection algorithm used at IHEC). Small peaks, instead, are more appropriate for quantification.

With the aim of quantifying the number of peaks in the histone modification signals, we designed our algorithm to identify small peaks. Let 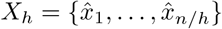 be a transformation of the profile *X* at resolution *h* (here 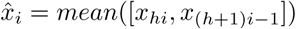), and let *R*(*X*_*h*_) = {*r*_1_, …, *r*_*m*_} (*m ≤ n/h*) be a compact representation of *X*_*h*_, where consecutive pairs 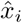 and 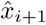 are merged if 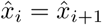. An element *r*_*i*_ is eligible as a peak if it satisfies, at least, the conditions: *r*_*i*_ *> r*_*i-*1_ and *r*_*i*_ *> r*_*i*+1_. We observe that the representation of a histone modification profile with *R*() allows to consider as candidate peaks all the maxima, independently on their width. However, in order for a point *r*_*i*_ to be a real peak, we need to verify also the steepness of the signal increase, and to compare *r*_*i*_ with its background. To this end, we partition the profile *R*() into intervals of the form *I*(*r*_*i*_) = [*α, β*], such that *r*_*α-*1_ = 0, *r*_*β*+1_ = 0, and *r*_*j*_ ≠ 0 ∀*j ∈*[*α < i < β*]. Finally, we compute the z–score of the peak intensity as:

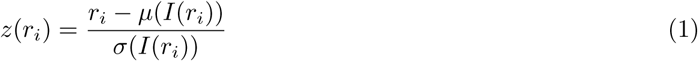

where *µ*() denotes the mean, and *σ*() is the standard deviation over the interval *I*(). The z–score defined in Eq. 1 is independent on the scale of the histone modification signal, and has the advantage of being interpretable as a sort of fold change. Consequently, we define a peak as a genomic locus where the score *z*() is higher than a user defined threshold (set to 2 in our experiments).

### 2.2 Normalization

After calling, we proceeded with counting the number of peaks for each gene. Peak counting, as many other quantification tasks from NGS data, is influenced by the sequencing depth. Indeed, in order for a peak to be individuated, it needs to be endowed with enough supporting reads. This generally happens easily with strong signals, while it requires high coverage for weaker signals. Counts Per Million (CPM) and Reads Per Kilobase per Million [13] (RPKM) are two widespread normalization methods used in the field of RNA–seq to mitigate the effect of sequencing productivity. Both methods leverage on the acceptable assumption that the overall amount of signals (in our case, peaks) per sample is roughly constant.

The main difference between these two methods is that RPKM supposes also that the molar concentration of RNA is constant and, consequently, the number of reads per gene is proportional to gene length. In our context, CPM and RPKM assume a somehow different semantics. CPM is an absolute measure of concentration of histone modification peaks on the gene, and it does not make any assumption on its distribution across the gene. RPKM, in contrast, is a relative measure. The rationale, in this second case, is that in order to produce a phenotype, the presence of a high concentration of a given histone modification is not sufficient itself, but it has to be spread along the gene.

Due to the lack of evidence in support of either a model based on the absolute concentration of histone modifications, or one based on a relative concentration, we tested both normalization methods in our experiments.

### 2.3 Cell type expression profiles

The IHEC histone modification data portal makes a variable number of different samples available for a given cell type. We exploited this redundancy by creating a signal–specific histone modification profile for each cell type, in order to mitigate the effect of intrinsic per sample variability. In short, we first computed a cell type profile averaging the contributions of the corresponding samples, and then used feature selection to retain only significant genes.

Similarly to what happens in the study of gene expression, it is sensible to assume that most genes do not contribute to a phenotype of interest, since they have either constant or no expression. Although apparently not interesting, these genes play a relevant role when computing distances between sample profiles. Indeed, they add constant factors to the distance computation with the effect of stretching distances between samples, and this causes the ratio between the two closest elements and the two farthest elements to tend to 1. Filtering out these genes would therefore mitigate this stretching effect without altering the ordering of pairwise distances.

Setting a cutoff threshold could potentially include/exclude genes with similar histone modification profiles only on the basis of a negligible distance from the threshold. We solved this problem by clustering genes and using the centroids as representatives of all cluster members. This ensures that genes with a similar profile are both either filtered or retained. We used a conservative threshold by requiring the maximum value of the centroid on a sample to be higher than the lowest 10% of the expression interval. The k–means algorithm was applied, endowed with the Lloyd procedure described in [11] (for the implementation we used the R [16] function `kmeans`), to select initial centroids to perform clustering. We employed a large value for *k* (as large as *k* = 50) to ensure high within–cluster homogeneity.

### 2.4 Similarity network fusion

The first tool that has been used to perform the experimental analysis presented in this paper is called *Similarity Network Fusion* (SNF) [21]^2^. In brief, given multiple similarity networks obtained from different data types referring to the same set of units, SNF integrates them into a unique similarity network through a cross diffusion process (CrDP) [20] in such a way that two types of links are promoted [21]: (i) Strong links, that are not necessarily present in all the networks, and (ii) links that are shared by all the networks. The output similarity network can be viewed as a combination of the significant features coming from different data types. Therefore, the fused network can give information on how multiple variables determine similarities among the studied objects.

### 2.5 Hypothesis testing scheme: Hypothesis vs. maximum cut

In this section, we introduce a quantitative method for verifying the plausibility of the differentiation between myeloid and lymphoid lineages, which characterizes the first hierarchical subdivision in the haematopoietic tree. The goodness of such hypothesis is evaluated by applying two different cuts to networks of distances among cell types.

An outline of the hypothesis testing model applied to the experiments is represented in Figure 2.

**Figure 2:**
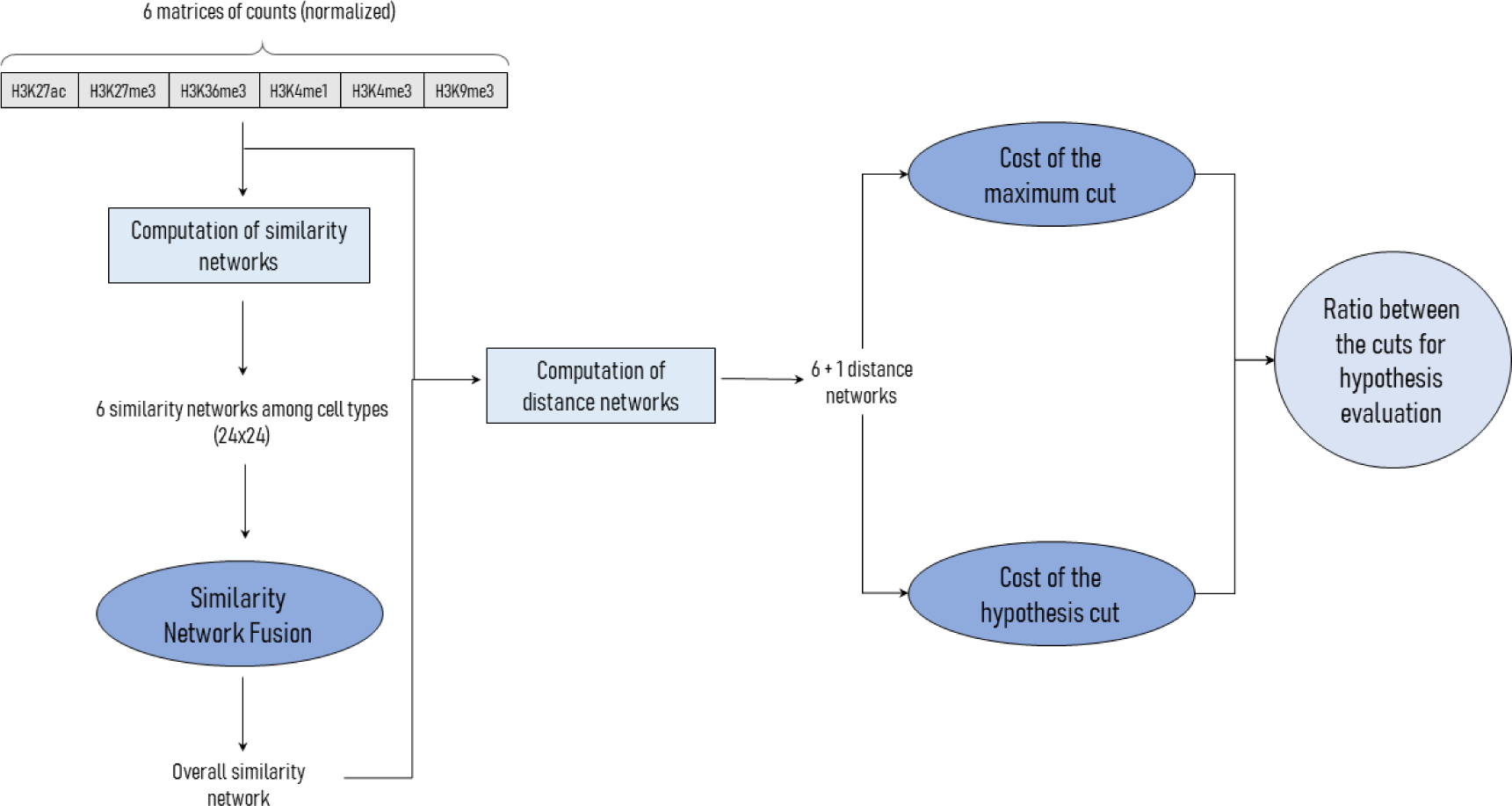
Model outline for testing the differentiation hypothesis between myeloid and lymphoid lineages in haematopoiesis. After the fusion process, the fused similarity matrix is transformed into a dissimilarity network. Then, the comparison between the cost of the hypothesis cut and of the maximum cut is applied both to the single data type dissimilarity networks and to the fused network.

Let *X* = {*x*_1_, …, *x*_*n*_} ⊆ ℝ*m* denote the experimental units, which are represented by *m* dimensional vectors. Moreover, let *d*: *X* × *X* →ℝ denote a distance or a dissimilarity measure on *X*. The computation of the distance/dissimilarity between pairs of experimental units allows to define the symmetric matrix *D* = (*d*(*x*_*i*_, *x*_*j*_))_*i,j*=1,…,*n*_, which constitutes the adjacency matrix of the distance/dissimilarity network of the considered units.

Suppose now that we want to evaluate the plausibility of a classification of the experimental units into two distinct classes, namely, *C*1 and *C*2. To this aim, each node of the dissimilarity network is labeled according to the class it belongs to. The problem of verifying how good the distinction between the two classes is reduces to quantifying the cost of bi–partitioning the network into the two sets corresponding to the classes, and to comparing it with the cost of the best cut.

A quantification of the cost of the hypothesis can be obtained by bi–partitioning the graph separating the two classes, and then computing the cost of the cut as the sum of the eliminated edges. The cost of the best cut can be obtained by computing the cost of the maximum cut of the dissimilarity network. Indeed, a max–cut algorithm applied to a dissimilarity network individuates a bi–partition where the dissimilarity within each set of the bi–partition is minimized, and the dissimilarity between the two sets is maximized. Although the problem of finding the maximum cut of a graph is known to be *NP* –complete [10], there are proposals of heuristic solutions, such as that given in [2].

The procedure presented in [2] — called *Greedy Cut algorithm* — starts by randomly selecting an edge, and by using its endpoints to initialize the sets of the bi–partition. Then, all the other vertices are associated to one of the two sets on the basis of an attraction function. The idea is to associate a vertex to the nearest set of the bi–partition. Once all the vertices are organized into a bi–partition, the cost of the cut is computed and compared with that at the previous iteration. The procedure is iterated enough to ensure that all the edges are considered [2].

## 3 Experiments

### 3.1 Dataset

For our tests, we used a collection of samples of cells of multiple stages in the haematopoietic cell lineages. Data were taken from the blueprint consortium^3^, available through the *International Human Epigenome Consortium* (IHEC) [9] data portal.

The database consists of healthy and diseased genome–wide profiles of several samples for each of the 6 marks on the Histone H3 identified by the Roadmap Epigenome Mapping Centers ^4^: the mono and tri– methylation of lysine 4 (H3K4me1 and H3K4me3), the tri–methylation of lysine 27 and 36 (H3K27me3 and H3K36me3) and the acetylation of lysine 27 (H3K27ac). Being interested to the physiological process of haematopoietic differentiation, we limited our experiments only to healthy samples.

Due to a limitation of the similarity network fusion method, which requires all the single data type networks to have the same nodes, we narrowed our experiment only to cell types for which all the histone modification marks were available. At the end of this filtering step, we obtained a dataset consisting of 24 distinct cell types, half belonging to the myeloid lineage and half to the lymphoid one (see Table 1 for details on the cell types considered and on the number of samples available).

**Table 1:**
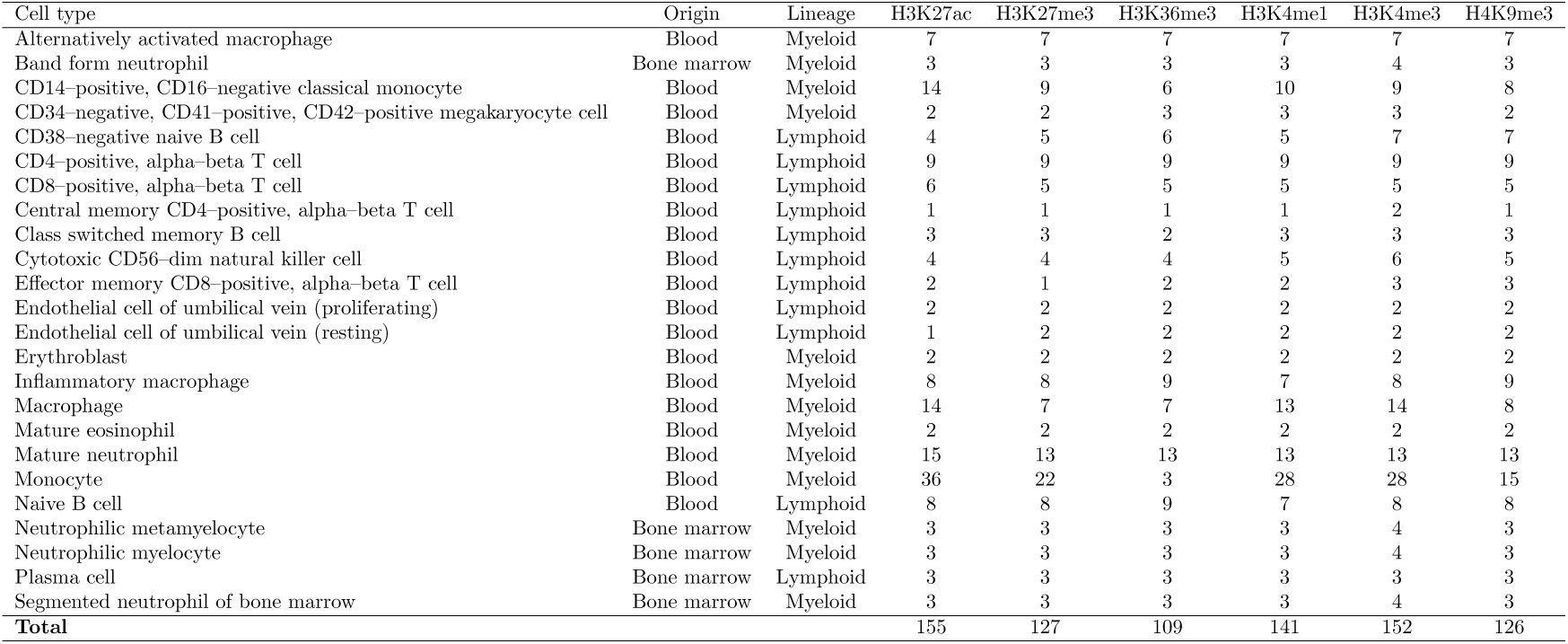
Origin, lineage, and number of samples for each cell type and histone modification.

### 3.2 Histone signal distribution

A first question we attempted to address is whether histone modification marks behave somehow similarly to gene expression. As most genes have constant or no expression (and only a limited number of genes have a noticeable one), we investigate the possibility that a (relatively) high signal intensity of a histone modification is registered only in a fraction of genes.

To answer this question, in table 2 we report the number of genes retained after feature selection. Experiments were conducted in the R environment [16] and, in particular, CPM and RPKM normalization were realized, respectively, with the R functions `cpm` and `rpkm` from the **edgeR** package [17]. Results show that, out of 21, 987 quantified genes, only a fraction of them is active. In particular, using RPKM, thus requiring the signal intensity to be proportional to the gene length, the number of active genes is rather little. Interestingly, this seems to be independent of the type of histone modification mark.

**Table 2:** Number of retained genes after feature selection either after CPM or RPKM normalization.

| Modification | CPM | RPKM | INTERSECTION |
| --- | --- | --- | --- |
| H3K27ac | 5.655 | 481 | 340 |
| H3K27me3 | 5.294 | 235 | 184 |
| H3K36me3 | 6.062 | 369 | 264 |
| H3K4me1 | 7.309 | 248 | 206 |
| H3K4me3 | 5.627 | 235 | 189 |
| H3K9me3 | 5.295 | 383 | 280 |

Comparing CPM and RPKM, the latter normalization filters much more genes. This result matches expectations, since large genes can have enough histone modification marks to be highlighted with CPM, but with not sufficient concentration to emerge with RPKM. Inspecting the intersection of the two sets of genes, however, we found that there is a portion of genes that passes the RPKM–based filtering, but not the CPM–based one. An in–depth examination of these genes shows that they have little peaks, and they emerge only because of their short length.

### 3.3 Myeloid/lymphoid differentiation evaluation

In this section, we report the results about the use of our model to assess the hypothesis of the differentiation on the haematopoietic lineage tree at myeloid/lymphoid level. As explained in Section 2.5, the proposed method for testing the hypothesis starts from the construction of dissimilarity networks between cell types. The dissimilarity networks were obtained in two different ways corresponding, respectively, to single histone modification profiles and to the fused similarity network. For the single modification profiles, the dissimilarity between pairs of cell types was computed as the squared Euclidean distance normalized in the range [0, 1]. For the fused similarity network resulting from the SNF procedure, instead, dissimilarities were computed by starting directly from the similarity network. More precisely, we first computed the z-–score of each similarity value, so that similarities were centered with respect to the mean value. Then, we inverted the z–scores with respect to the mean, thus obtaining a dissimilarity network of cell types.

In this approach, if a given hypothesis is supported by data, partitioning the network on the basis of it should cause the removal of only highly weighted edges. We can thus use the cost of the cut induced by this partitioning as a measure of how much the tested hypothesis is supported by data.

The coarse cost of the cut is difficult to interpret, due to its dependence on the dissimilarity function. In order to get rid of scale problems, we converted it into a scale-free score according to the following equation:

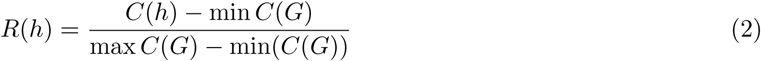

where *G* is the considered network, *h* is the hypothesis cut, and the function *C*() denotes the cost of a graph cut. The cost of the maximum cut was computed by implementing in the R environment the Greedy Cut Algorithm described in Section 2.5. The cost of the minimum cut was obtained by using the R function `min cut` from the **igraph** package, which implements the minimum cut of a graph following the algorithm proposed in [18].

The score in Eq. (2) ranges between 0 and 1 assuming maximum value 1 when the cost of the hypothesis equals that of the maximum cut on *G*.

Figure 3 shows a comparison of the hypothesis score for the two normalization strategies (CPM and RPKM) both for the SNF network and for the individual histone modification networks.

**Figure 3:**
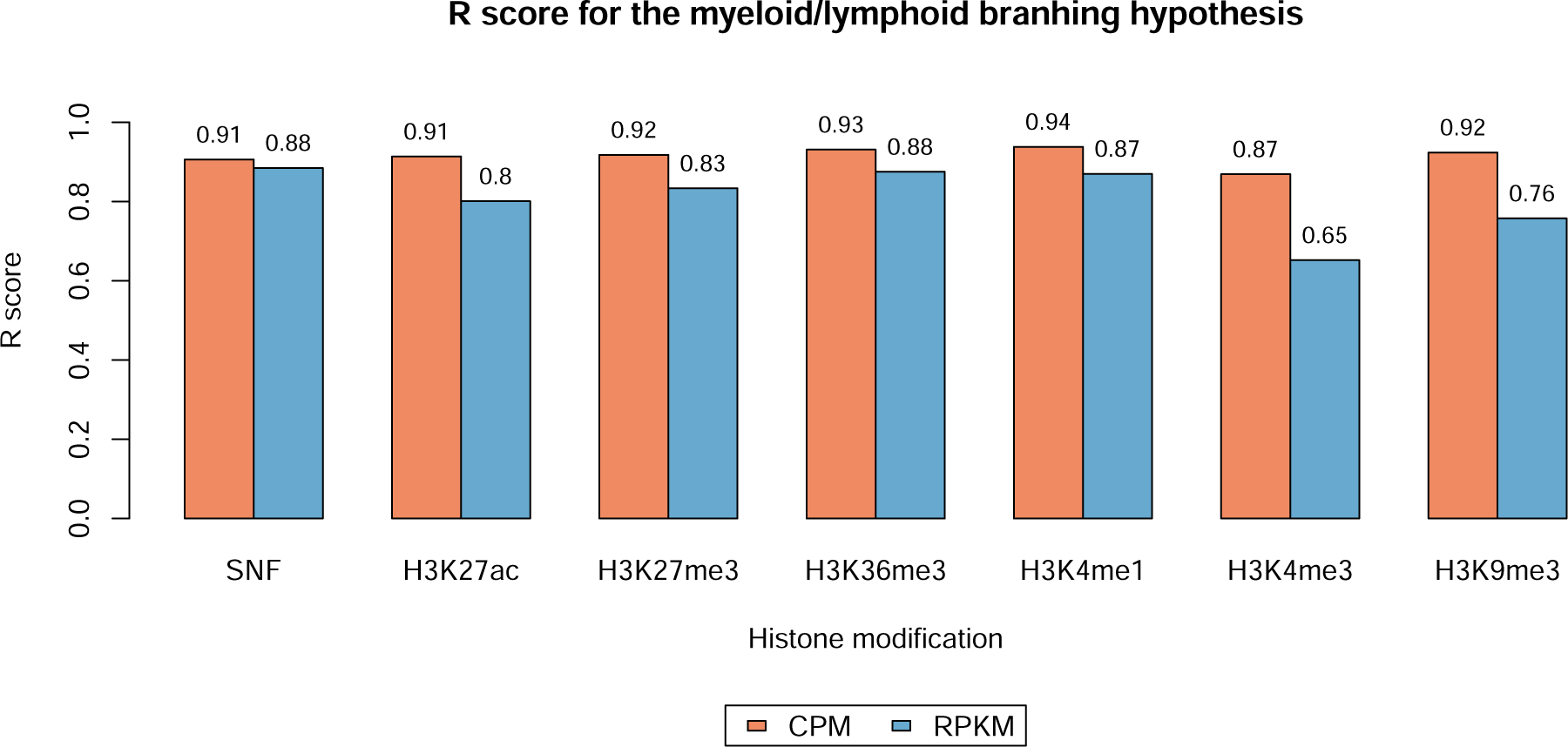
Barplot of the results of the hypothesis testing scheme for each modification network, and for the SNF network.

At a glance, we observe that scores obtained on CPM based normalization are consistently higher than those obtained with RPKM. This suggests that, in order to trigger a certain phenotype, histone modifications do not need to be uniformly spread along the gene, but it is enough to have their presence in sufficient concentration. Comparing the histone modifications, Figure 3 shows that all the types of histone modification almost equally contribute to the haematopoietic branch at this level. This is confirmed also by the analysis of the SNF network. Finally, although scores as high as those shown in Figure 3 agree with the classical myeloid/lymphoid branch depicted in Figure 1, they suggest that this model cannot be the only mechanism for haematopoietic blood cell differentiation.

## 4 Discussion

Histone modifications are complex signals which are not completely understood yet. It is known that an increase/decrease in the concentration of such signals has an impact on the gene expression. In addition, from [3], we know that the presence of large peaks in the signal wave is associated to loci involved in a histone modification. In our experiments, a step forward was made, consisting in the investigation of the possibility of using peaks at high resolution to quantify per gene histone modifications. Following this idea, we studied the distribution of high resolution peaks across the genome, with the result that such peaks behave similarly to gene expression profiles. More specifically, it can be observed, similarly to what happens for gene expression, that only a small fraction of genes have a noticeable or high signal intensity. Moreover, we verified that per gene unbalanced concentrations of peaks can be associated to different phenotypes.

Besides sequencing productivity, the number of peaks per gene can also be greatly influenced by gene length, when working at high resolution. Nonetheless, performing differential analysis tasks, gene length becomes a constant and, thus, normalizing counts could be not necessary. In our case, however, the adoption of a normalization based on gene length can be viewed as a change in the interpretation of the quantification of a histone modification: a histone modification is thought of as a signal distribution, rather than a measure of concentration. With the aim of clarifying which one of the above interpretations better reflects the mechanisms underneath histone modifications, we tested two normalization methods: CPM and RPKM^5^. CPM corresponds to a measure of concentration, namely, a model in which differences on the phenotype are triggered when a sufficiently high amount of modifications occurs across the gene, independently on its distribution. This model is coherent with the idea that histone modifications have the mere role of starting/stopping transcription. On the other hand, RPKM is a measure of distribution. In this case, high quantification values for large genes are achieved only when the peaks are distributed across the gene. In this case, the role of histone modifications is supposed to be that of keeping the entire sequence of the gene accessible/hidden, in order to facilitate/prevent transcription.

Results in Table 2 show that, similarly to the expression of genes, in most cases the signal (i.e. the number of peaks) is almost absent. Not surprisingly, this is particularly evident using the RPKM normalization. Indeed, long genes could have a number of peaks high enough to overcome the threshold for CPM, but not for RPKM. However, as the intersection of the relevant genes with CPM and with RPKM shows, the opposite phenomenon is also present: there are short genes whose concentration of peaks is not sufficiently high to pass the CPM filtering, but characterized by a signal distribution allowing to pass the RPKM one. An in–depth inspection of the results in Table 2 also reveals that the number of active genes is quite constant, regardless of the histone modification type. Although a precise biological interpretation of this fact would require further investigations, we could speculate about the fact that none of the histone modification signals seems to have a dominant role in the regulation of gene expression, but rather the displayed phenotype is the result of the combination of the individual contributions. For example, in imprinted genes both the open chromatin mark H3K4me3 and the compacted chromatin mark H3K9me3 are present at the promoter site [12].

Based on this assumption, we used the well–known SNF method [21] to integrate all the histone modification signals into a unified model. The resulting SNF network, shown in Figure 4, is a graph where nodes correspond to cell types and edges are weighted with a score proportional to the similarity between the nodes they connect.

**Figure 4:**
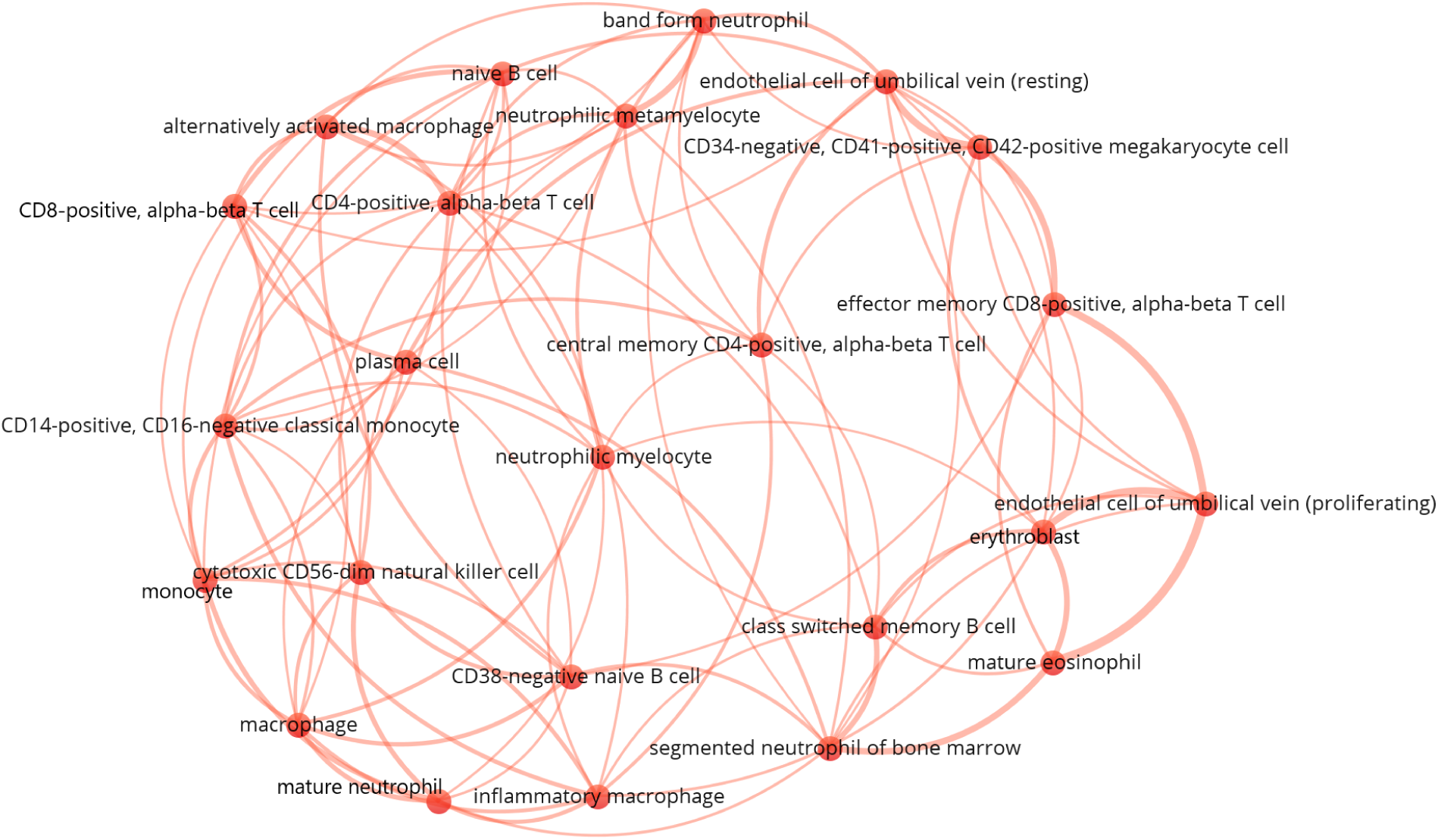
Similarity network of cell types obtained after the application of the SNF method. Here we chose to plot only the 100 strongest links, in order to reduce noise and to facilitate visualization. The plot is obtained with the VOSviewer software [19].

In order to apply our hypothesis testing model, we turned this graph into a dissimilarity network by computing the z–scores of the edge weights, and reversed the weights with respect to the mean weight. On the obtained dissimilarity network we tested the hypothesis of differentiation between the myeloid and the lymphoid lineages, i.e. the first step of the classical haematopoietic tree (see Figure 1). According to this model, the relationships between pairs of the same lineage should be stronger (hence dissimilarity scores should be lower) than relationships between the two lineages. Consequently, in the ideal case, the cost of the cut that partitions the SNF graph into the two clusters corresponding to the two lineages (one with the myeloid cells and one with the lymphoid ones) should be maximum. Results reported in Figure 3 show that this is almost the case, thus indicating that the classical myeloid/lymphoid differentiation branching is a reasonable approximation of the real haematopoietic process. However, the same results leave enough room to conclude that this model in not accurate enough to capture the complexity of haematopoiesis. Interestingly, narrowing to the single histone modification graphs, comparing the myeloid/lymphoid cut with the maximum cut, we found that all the signals approximate the classical model with comparable scores. This, again, can be considered a confirmation of the hypothesis that all the histone modification marks cooperate to the development of the displayed phenotype.

Overall, experiments have proved that histone modification marks can be quantified using high resolution peaks. This quantification behaves similarly to gene expression, with only few active genes at the same time. Moreover, experiments on a population involving 24 cell types belonging to the haematopoietic tree have shown that there is a causal relationship between a given phenotype and a profile of the modification marks. This opens to exploiting differential analysis to identify genes involved in a phenotype of interest.

## 5 Conclusions

Histone modifications are complex signals which regulate gene expression by modifying the tridimensional structure of the genome and, in turn, making genes more/less accessible for transcription. The complexity of these signals makes their mining very difficult.

In this paper, we show that counting peaks at high resolution (as high as few bps) can constitute a reasonable approach to build per-gene profiles of histone modification marks. The experimental analysis of signals from six histone modifications belonging to 24 cell types highlights that these profiles follow a distribution which is similar to that of the expression of genes. The meaningfulness of the peak-based histone profiles is validated by computationally assessing the classical lympoid/myeloid branch at the first level of the haematopoietic tree.

Our experiments confirm that the model fairly approximates the real haematopoietic process, although suggesting that it does not completely capture its complexity. Besides the contribution on the specific topic of haematopoiesis, our work constitutes an advance in the understanding of epigenetics by providing a framework to analyze histone modification data, as well as a methodology to validate and compare different hypotheses on a population of cell types.

Finally, the distribution of the signal of histone modification profiles enables using standard differential expression techniques to identify genes whose modifications are involved in a given phenotype.

## Acknowledgement

The authors would like to thank Prof. Pietro Liò for his useful advises and suggestions.

1 http://www.roadmapepigenomics.org/

2 The software can be downloaded in the *R* or *MATLAB* version at http://compbio.cs.toronto.edu/SNF/SNF/Software.html.

3 https://www.blueprint-epigenome.eu/

4 http://www.roadmapepigenomics.org/

5 Notice that, in our case, we substitute the number of reads with the overall number of peaks.

